# Tropical forest refuge in peripheral soils of slags in Caatinga

**DOI:** 10.1101/317685

**Authors:** José João Lelis Leal de Souza, Bartolomeu Israel de Souza, Rubens Teixeira de Queiroz, Rony Lopes Lunguinho, Joseilson Ramos de Medeiros, Eini Celly Morais Cardoso, Pedro André de Melo e Silva, Helder Cavalcante de Oliveira

## Abstract

The influence of environmental factors on the structure and composition of plant communities in the Caatinga is complex and poorly explored. Spatial variation of biodiversity in Caatinga is poorly know and strictly attributed to climatic conditions. We investigated the influence of slags on floristic composition and structure of a shrubby-arboreal community in one of the driest region in Brazil. Chemical and physical analyses of soils were performed in samples of seven plots from savannic formations and from forest formations. Vegetation was characterized floristically and structurally in all plots. Habitats were structurally distinct, and diversity differed between peripheral and non-peripheral areas of slags. Nine of the ninety-seven species identified are reported to (sub)humid biomes. Soils are dominantly shallow, eutrophic and sandy loam. However, soils in the periphery of slags are more developed once paludization, melanization and bioturbation were verified. Our results suggest that soil fertility did not influence vegetable cover in Caatinga. The cover of plant species considered exclusive of (sub)humid biomes in Brazil extends beyond highlands in the semiarid, associated with high soil organic carbon content and water retention capacity of more developed soils than the typical of the Caatinga.

## Introduction

Dryland biomes cover 41.5 % of the Earth’s surface, with seasonally dry tropical forests occupying approximately 1,327 million hectares within this area [1]. The largest area of continuous dry tropical rainforests in Latin America occurs in Brazil, with approximately 800.00 km^2^. Popularly called as Caatinga (Mata Branca), most trees and shrubs lose their leaves in the dry season, which can last up to nine months a year in certain regions [2].

Caatinga is a Brazilian exclusive biome composed by different plant formations [3] and its diversity is considered one of the most biodiverse among semiarid biomes [4,5]. Among the several ecosystems that exist in the Caatinga, granitic intrusions in the form of inselbergs, massive, slags (lajedos in portuguese) and bornhardts cover 15 % of the total area of the northern Brazilian Northeast [6].

Low developed, shallow and eutrophic soils are dominant in Caatinga due to tropical dry climate. However, high local diversity of soils has been attributed to variance of lithology and, or, relief by recent studies [7]. Understood soil diversity is a relevant task, especially considering that plant distribution is conditionate by soil properties [8,9]. Soil influences plant physiognomies in Brazilian Savanna (Cerrado) [10], Restinga forest [11], Amazonia rainforest [12,13], Tundra [14] and other biomes around the world [15,16]. However, species diversity in Caatinga is poorly know [17] and strictly attributed to climatic conditions [18–20].

The purpose of this study was to investigate a vegetation gradient at preserved slabs in Caatinga, and to describe the main soil-vegetation relationships and the community structure. Thus, we hypothesized that variation in vegetation structure and composition is related to soil chemical and physical attributes.

## Material and methods

### Study area

The study area encompasses the occidental and oriental Cariri microregions, Paraíba state (Fig 1). They have a total area of approximately 11,225.736 km^2^ and particular landscape diversity. Altitudes vary between 400 m, in the Paraíba valley, and 700 m, in highlands associated with resistant rocks and horst–graben systems. The mean precipitation of 350 mm year-1, potential evapotranspiration four times higher than precipitation and mean temperature of 27 °C highlights Cariri as one of the driest regions of Brazil [21].

**Fig 1.**
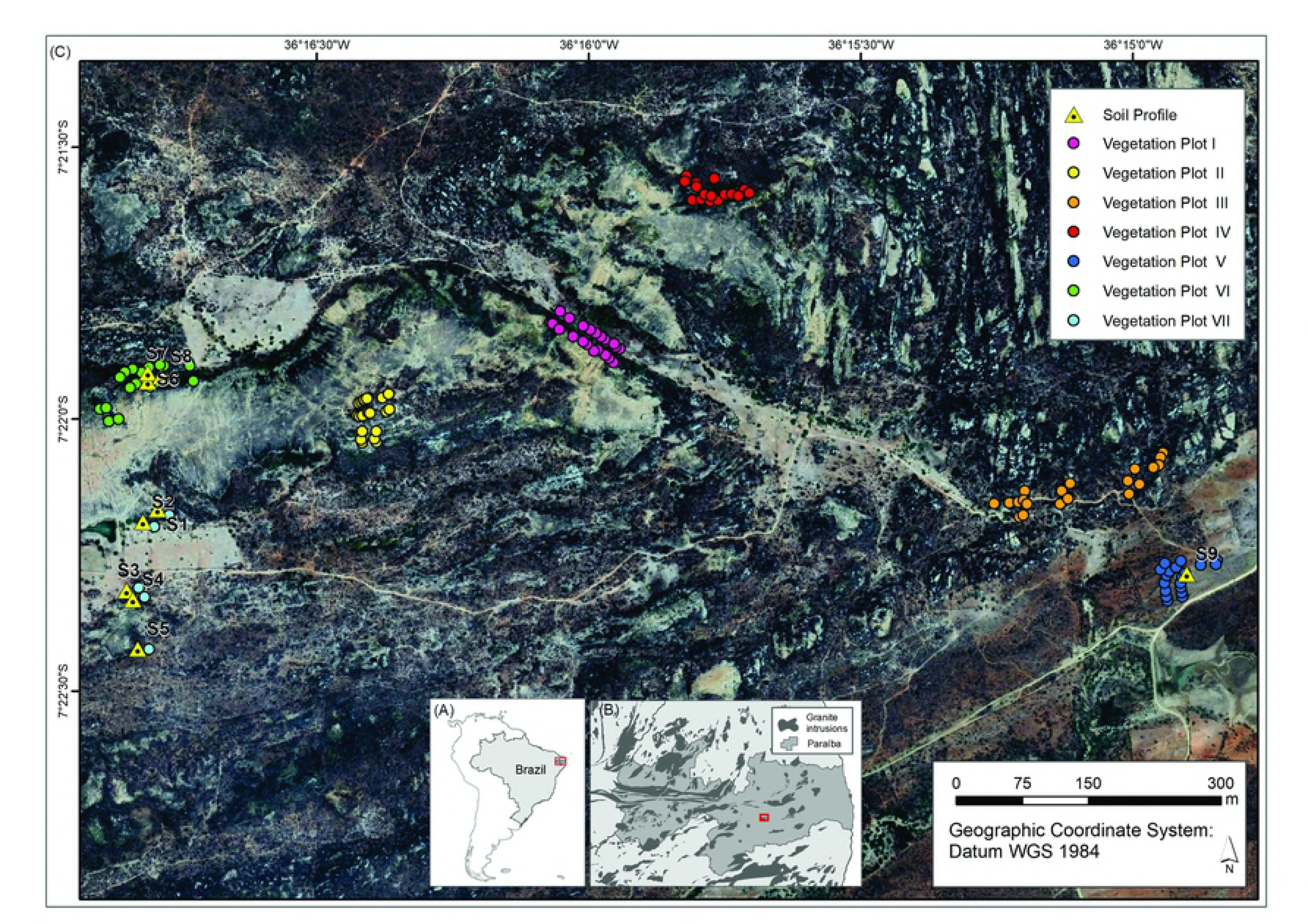
Paraíba State location (A), study area location (B) and sample distribution (C).

Cariri microregions are inserted in the Borborema province, characterized by the extensive erosive surfaces denominated the *sertanejas* depressions developed on Meso- and Neoproterozoic gneiss and schist [22]. Inselbergs, massives and slabs developed on Neoproterozoic granites emerge due to differential weathering by mineralogical, petrographic or fracture pre-arrangement [6].

In Cariri microregions shallow and incipient weathered soils dominate, with Entisols (Neossolos according to Brazilian System of Soil Classification) and Alfisols (Luvissolos) representing 43 % of the study area [23]. Plant physiognomy is described as dry deciduous forest dominated by Bromeliaceae Juss., Cactaceae Juss. and Fabaceae Lindl. and members of Poaceae Barnhart and Cyperaceae Juss. families [24].

### Data sampling and analysis

Sixty-five transects, 50 m × 2 m each one, were established in Environmental Protection Area of Cariri between January and November 2017, according to linear transect method for phanerophytes and camphids (LTMPC) [25]. The LTMPC was successfully adopted in previous studies in Caatinga [26]. Seven different vegetation plots (V.P.) were formed by grouping transects: a) V.P. on peripheral areas of slags: V.P. I, V.P. II, V.P. IV and V.P. VI; b) V.P. on a flat plain: V.P. III and V.P. V; c) V.P. on peripheral areas of slags and on flat plain: V.P. VII.

Height, the largest and the smallest diameters of individuals with diameter at breast height below 2 cm were measured. Individuals with diameter at breast height above 2 cm had their height and median crown radius measured. All live specimens were sampled and identified at field and consulting Brazilian Flora 2020 [27]. The APG IV classification system was adopted for the plant families [28].

The Jaccard similarity index was calculated to express the similarity between plots based on the number of common species. The resulting floristic similarity matrix was used for cluster analysis by simple binding method and generation of a dendrogram in the PAST software [29].

### Soil sampling and analysis

Seven pedons were taken, described and classified according to Soil Taxonomy (Soil Survey Staff, 2014) and Brazilian Soil Classification System [30] to represent soil diversity of plots. Soil samples were collected from the surface down to the lithic contact at each pedon. For deeper soils a 200 cm control section was used. Because the absence of temperature soil data in Brazilian semiarid, we classified the moisture regime of soils according to previously studies [31].

Samples were air dried and sieved through a 2 mm sieve prior to physic and chemical analyzes according to methods established for tropical soils [32]. Coarse and fine sand, silt and clay were determined by the sieve-pipette method after dispersion with 0.1 M NaOH. Soil pH was measured with a glass electrode in a 1:2.5 suspension v/v soil and deionized water (H_2_O pH) and 1 M KCl solution (KCl pH). Delta pH (ΔpH) was calculated by KCl pH minus H_2_O pH. The potential acidity (H + Al) was extracted by 1 M ammonium acetate solution at pH 7. The content of exchangeable Ca^2+^, Mg^2+^ and Al^3+^ were determined in a 1 M KCl extract. Exchangeable K^+^ and Na^+^ were determined after Melhich-1 extraction. From these results, the sum of bases (SB), base saturation (V), aluminum saturation (m), equivalent cation exchange capacity (ECEC), total cation exchange capacity (CEC) and Na saturation (ISNA) were calculated.

The available phosphorus content (P_M_) was determined by a Mehlich-1 extraction solution. Total organic carbon (C) was determined by wet combustion [33]. Total nitrogen (N) was determined by Kjeldahl method and titration [34]. Carbon to nitrogen ratio (C/N) was calculated on the mass basis. The P adsorption capacity of the soil was determined after stirring for 1 hour with 2.5 g of soil in 0.01 M CaCl_2_ containing 60 mg of P L^-1^. The suspension was filtered and the remaining P in solution (P_REM_) was determined by photocolorimetry (Alvarez et al., 2000).

Bulk density, particle density and micropores (pores with diameter below 0.5 mm) were determined in undisturbed soil samples collected by volumetric rings [34]. Total porosity and macropores (pores with diameter above 0.5 mm) were calculated from these results. Total organic carbon and nitrogen stocks were calculated by using the total organic content, sampling depth and bulk density: Stock (Mg ha^-1^) = bulk density (kg dm^-3^) x total organic carbon (or total nitrogen) content (%) x depth (cm).

Descriptive statistics were calculated for pedons according to its proximity of slags. Furthermore, Pearson correlation between soil properties was calculated. Principal Component Analysis (PCA) was performed to elucidate correlation between variables. Prior to PCA, analytical data of each horizon was logarithmic transformed and standardized to provide a normal distribution [35].

## Results

A total of 2,563 specimens were identified in all transects (Table 1), exceeding the recommendation for phytosociological surveys aiming to characterize Caatinga [36]. Six hundred and thirty-seven specimens (24.9 % of total) composes the arboreal stratum. The richness of species is dominated by Fabaceae (14 species), followed by Euphorbiaceae Juss. (7 spp.), Myrtaceae Juss. (4 spp.), Anacardiaceae R. Br. (4 spp.) and Rubiaceae Juss. (3 spp.) familes (S1 Table).

**Table 1.** Specimens distribution by strata.

| Height (m)/ strata | Number of specimens | Relative frequency (%) |
| --- | --- | --- |
| 0 - 0.3/ herbaceous to sub shrubby | 242 | 9.44 |
| > 0.3 - 0.6/ sub shrubby | 317 | 12.37 |
| > 0.6 - 1.5/ shrubby | 709 | 27.66 |
| > 1.5 - 3/ shrubby to arboreal | 658 | 25.67 |
| > 3 – 5/ small tree | 310 | 12.10 |
| > 5 – 10/ medium tree | 238 | 9.29 |
| > 10 – 20/ large tree | 73 | 2.85 |
| > 20/ largest tree | 16 | 0.62 |
| All | 2,563 | 100 |

Nine of the ninety-seven species identified are reported to other biomes (S1 Table): *Allophylus quercifolius* (Mart.) Radlk. is commonly associated to Amazon rainforest, Atlantic forest and Brazilian Savanna; *Calyptranthes lucida* Mart. *ex* DC and *Vitex orinocensis* Kunth are associated to Amazon rainforest and Atlantic forest; *Chloroleucon tortum* (Mart.) Pittier *Myroxylon peruiferum* L. F. are associated to Atlantic forest and Brazilian Savanna; *Erythroxylum suberosum* A. St.-Hil. is associated to Amazon rainforest and Brazilian Savanna; *Hymenaea rubriflora* Ducke, *Libidibia ferrea var. leiostachya* (Benth.) L. P. Queiroz and *Pisonia ambigua* Heimerl are associated to Atlantic forest.

The highest number of specimens were identified in vegetation plots (V.P.) peripheral to slags: V.P. 6 (755 specimens or 29.4 % of total), V.P. 4 (440 or 17.2 %), V.P. 2 (393 or 15.3 %) and V.P. 1 (370 or 14.4 %) (S1 Table). The highest number of specimens of the species associated to wetter biomes also are reported in V.P. peripheral of slags, corresponding to approximately 2.8 % of specimens in each V.P.

The values of coefficient of similarity vary from 27 % to 54 % (Fig 2). Higher Jaccard index values were observed for comparations between V.P. II, V.P. IV and V.P. VI indicating that these plots share a high number of species in common.

**Fig 2.**
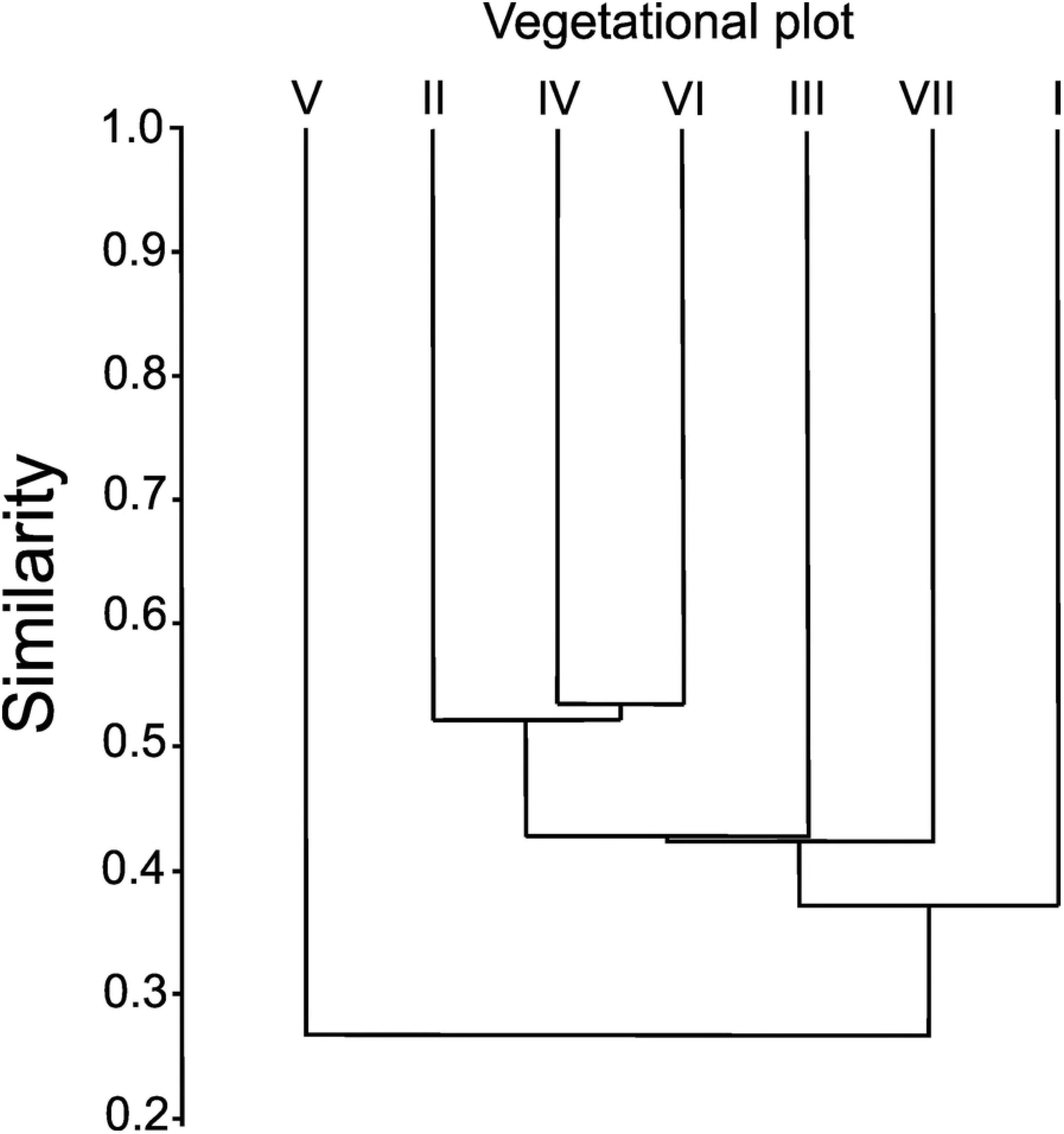
Similarity index between vegetational plots.

## Morphology and physic-chemical soil properties

Six of the nine collected soil profiles were classified as Entisols, which corresponds to the order of Neossolos in the Brazilian Soil Classification System [30]. Two Inceptsols were described: one Aridic Haplustepts with fragipan (Neossolo Regolítico) and the another one was identified as Oxic Haplustepts (Cambissolo Háplico). One Lithic Rhodustalfs (Luvissolo Crômico) were described to represent soil dominance of Brazilian semiarid (Table 2).

**Table 2.**
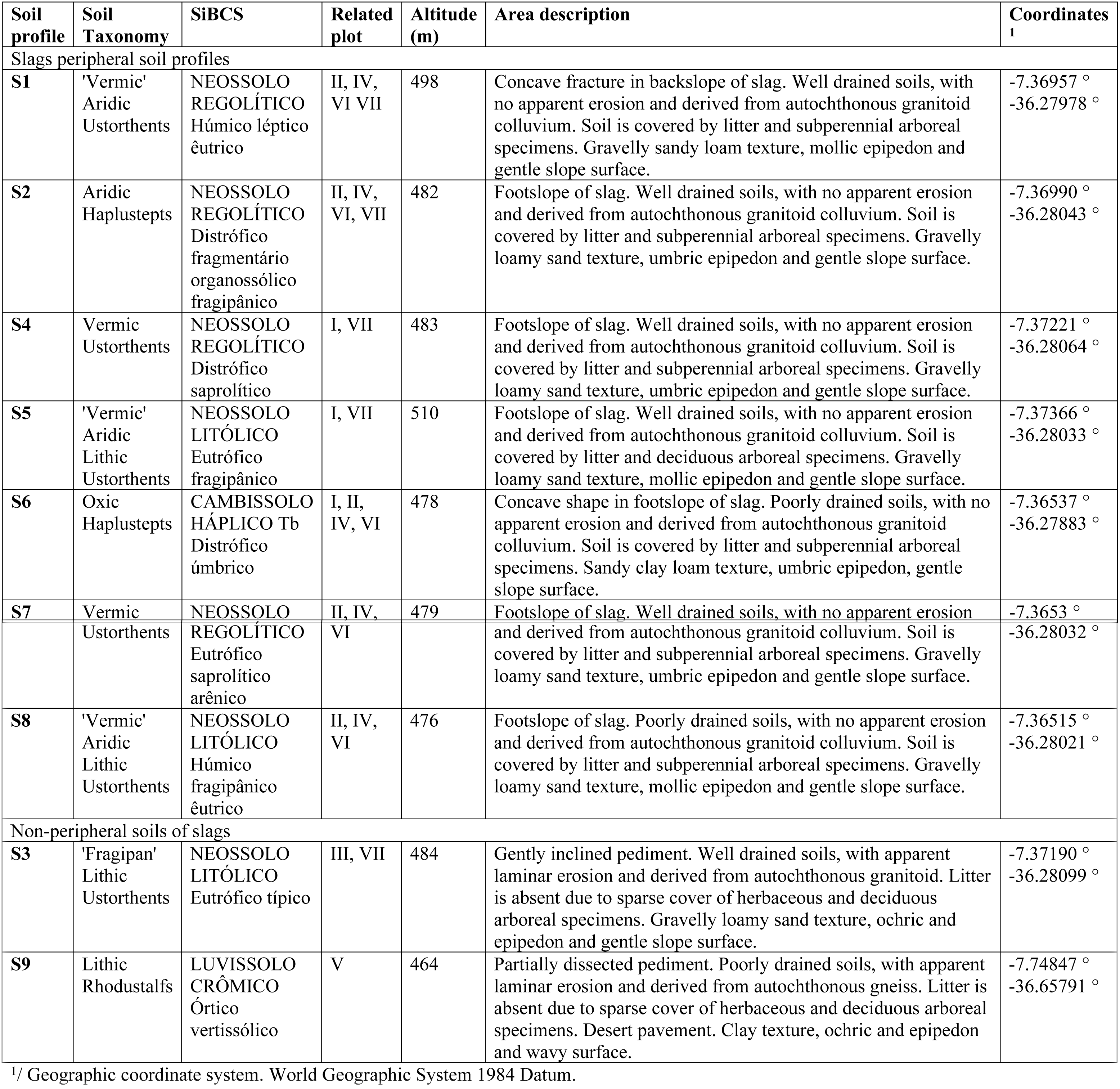
Area description and classification of soils.

| Soil profile | Soil Taxonomy | SiBCS | Related plot | Altitude (m) | Area description | Coordinates<sup>1</sup> |
| --- | --- | --- | --- | --- | --- | --- |
| Slags peripheral soil profiles |  |  |  |  |  |  |
| **S1** | 'Vermic'<br>Aridic<br>Ustorthents | NEOSSOLO<br>REGOLÍTICO<br>Húmico léptico<br>êutrico | II, IV,<br>VI VII | 498 | Concave fracture in backslope of slag. Well drained soils, with no apparent erosion and derived from autochthonous granitoid colluvium. Soil is covered by litter and subperennial arboreal specimens. Gravelly sandy loam texture, mollic epipedon and gentle slope surface. | -7.36957 °<br>-36.27978 ° |
| **S2** | Aridic<br>Haplustepts | NEOSSOLO<br>REGOLÍTICO<br>Distrófico<br>fragmentário<br>organossólico<br>fragipânico | II, IV,<br>VI, VII | 482 | Footslope of slag. Well drained soils, with no apparent erosion and derived from autochthonous granitoid colluvium. Soil is covered by litter and subperennial arboreal specimens. Gravelly loamy sand texture, umbric epipedon and gentle slope surface. | -7.36990 °<br>-36.28043 ° |
| **S4** | Vermic<br>Ustorthents | NEOSSOLO<br>REGOLÍTICO<br>Distrófico<br>saprolítico | I, VII | 483 | Footslope of slag. Well drained soils, with no apparent erosion and derived from autochthonous granitoid colluvium. Soil is covered by litter and subperennial arboreal specimens. Gravelly loamy sand texture, umbric epipedon and gentle slope surface. | -7.37221 °<br>-36.28064 ° |
| **S5** | 'Vermic'<br>Aridic<br>Lithic<br>Ustorthents | NEOSSOLO<br>LITÓLICO<br>Eutrófico<br>fragipânico | I, VII | 510 | Footslope of slag. Well drained soils, with no apparent erosion and derived from autochthonous granitoid colluvium. Soil is covered by litter and deciduous arboreal specimens. Gravelly loamy sand texture, mollic epipedon and gentle slope surface. | -7.37366 °<br>-36.28033 ° |
| **S6** | Oxic<br>Haplustepts | CAMBISSOLO<br>HÁPLICO Tb<br>Distrófico<br>úmbrico | I, II,<br>IV, VI | 478 | Concave shape in footslope of slag. Poorly drained soils, with no apparent erosion and derived from autochthonous granitoid colluvium. Soil is covered by litter and subperennial arboreal specimens. Sandy clay loam texture, umbric epipedon, gentle slope surface. | -7.36537 °<br>-36.27883 ° |
| **S7** | Vermic | NEOSSOLO | II, IV, | 479 | Footslope of slag. Well drained soils, with no apparent erosion | -7.3653 ° |
|  | Ustorthents | REGOLÍTICO<br>Eutrófico<br>saprolítico<br>arênico | VI |  | and derived from autochthonous granitoid colluvium. Soil is covered by litter and subperennial arboreal specimens. Gravelly loamy sand texture, umbric epipedon and gentle slope surface. | -36.28032 ° |
| **S8** | 'Vermic'<br>Aridic<br>Lithic<br>Ustorthents | NEOSSOLO<br>LITÓLICO<br>Húmico<br>fragipânico<br>êutrico | II, IV,<br>VI | 476 | Footslope of slag. Poorly drained soils, with no apparent erosion and derived from autochthonous granitoid colluvium. Soil is covered by litter and subperennial arboreal specimens. Gravelly loamy sand texture, mollic epipedon and gentle slope surface. | -7.36515 °<br>-36.28021 ° |
| Non-peripheral soils of slags |  |  |  |  |  |  |
| **S3** | 'Fragipan'<br>Lithic<br>Ustorthents | NEOSSOLO<br>LITÓLICO<br>Eutrófico típico | III, VII | 484 | Gently inclined pediment. Well drained soils, with apparent laminar erosion and derived from autochthonous granitoid. Litter is absent due to sparse cover of herbaceous and deciduous arboreal specimens. Gravelly loamy sand texture, ochric and epipedon and gentle slope surface. | -7.37190 °<br>-36.28099 ° |
| **S9** | Lithic<br>Rhodustalfs | LUVISSOLO<br>CRÔMICO<br>Órtico<br>vertissólico | V | 464 | Partially dissected pediment. Poorly drained soils, with apparent laminar erosion and derived from autochthonous gneiss. Litter is absent due to sparse cover of herbaceous and deciduous arboreal specimens. Desert pavement. Clay texture, ochric and epipedon and wavy surface. | -7.74847 °<br>-36.65791 ° |

<sup>1/</sup> Geographic coordinate system. World Geographic System 1984 Datum.

The depth of the soil varied between 20 cm and 100 cm with a mean of 72 cm (Table 3). Soils are shallow and transition between horizons was predominantly clear or abrupt, indicating that the parent material is low weathered. Structure is low to moderate developed, with sub angular blocky and single gran types dominantly. Worm hole sand filled animal burrows were observed in five Entisols, indicating widespread bioturbation. Soils without bioturbation were only observed in non-peripheral soils of slags.

**Table 3.**
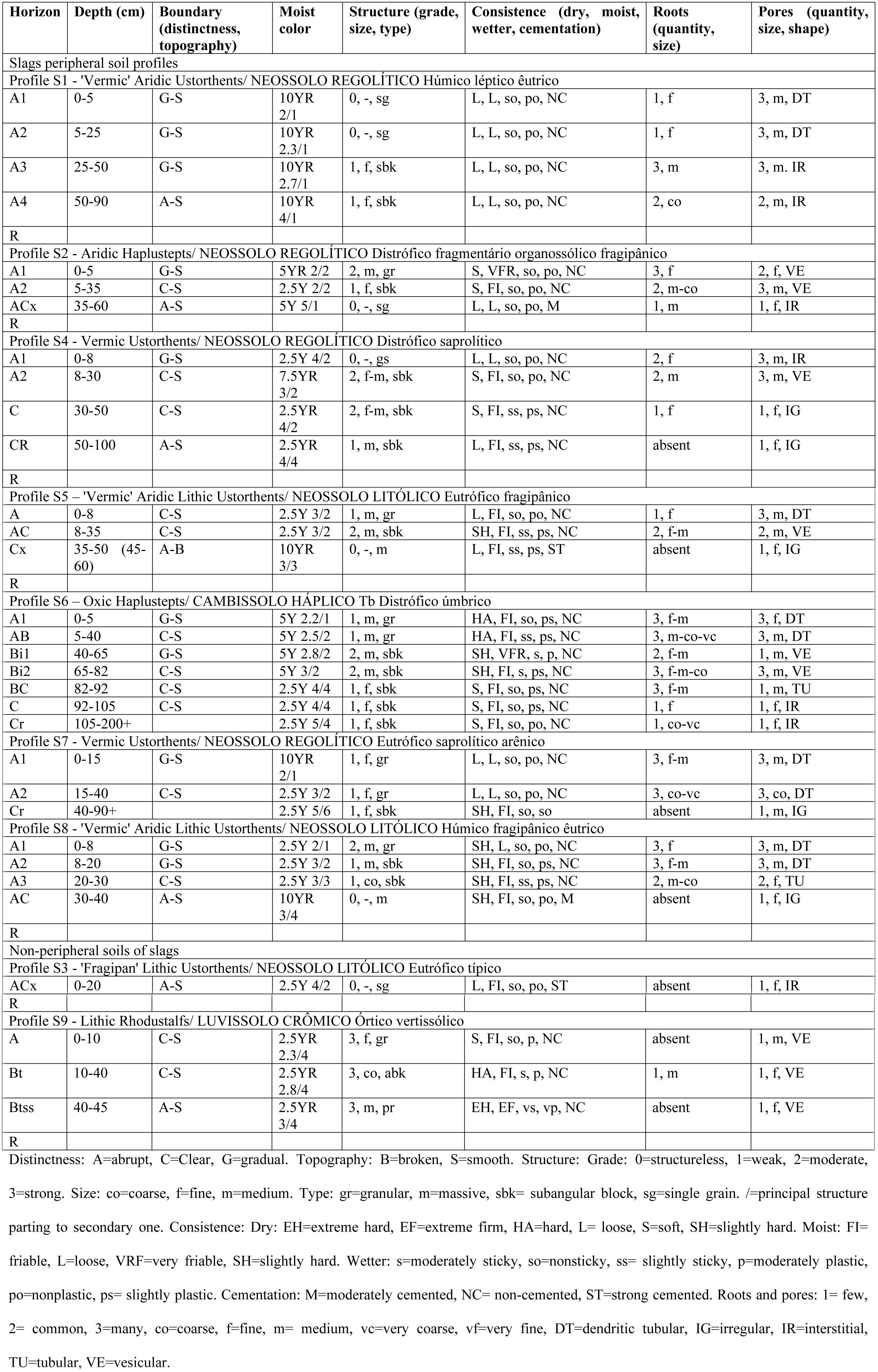
Morphological properties of soils.

Fragipan and fragic soil properties were observed in four soil profiles. Roots are denser in shallow layers and its growing are limited by lithic contact or fragipan. Medium and coarse roots occur below 30 cm of depth.

The soil color ranged from dark olive gray in cambic horizon to dark gray in umbric epipedon. Mollic and umbric epipedons described in slags peripheral soil profiles indicates paludization and melanization. Organic horizons described according to Brazilian Soil Classification System were presented as mineral horizons due to the higher limit of organic carbon in the Soil Taxonomy’s definition of organic soil material.

Texture is dominantly sandy loam and a slightly increase of clay content in deeper horizons is observed (Table 4). Feldspars were identified in the field as the main component of coarse fraction in all soils. Coarse sand is the main fraction of soil, once its mean content is 50 %. The deepest soil, Oxic Haplustepts, presented mean clay content until 2.4 times higher than the other soils.

**Table 4.**
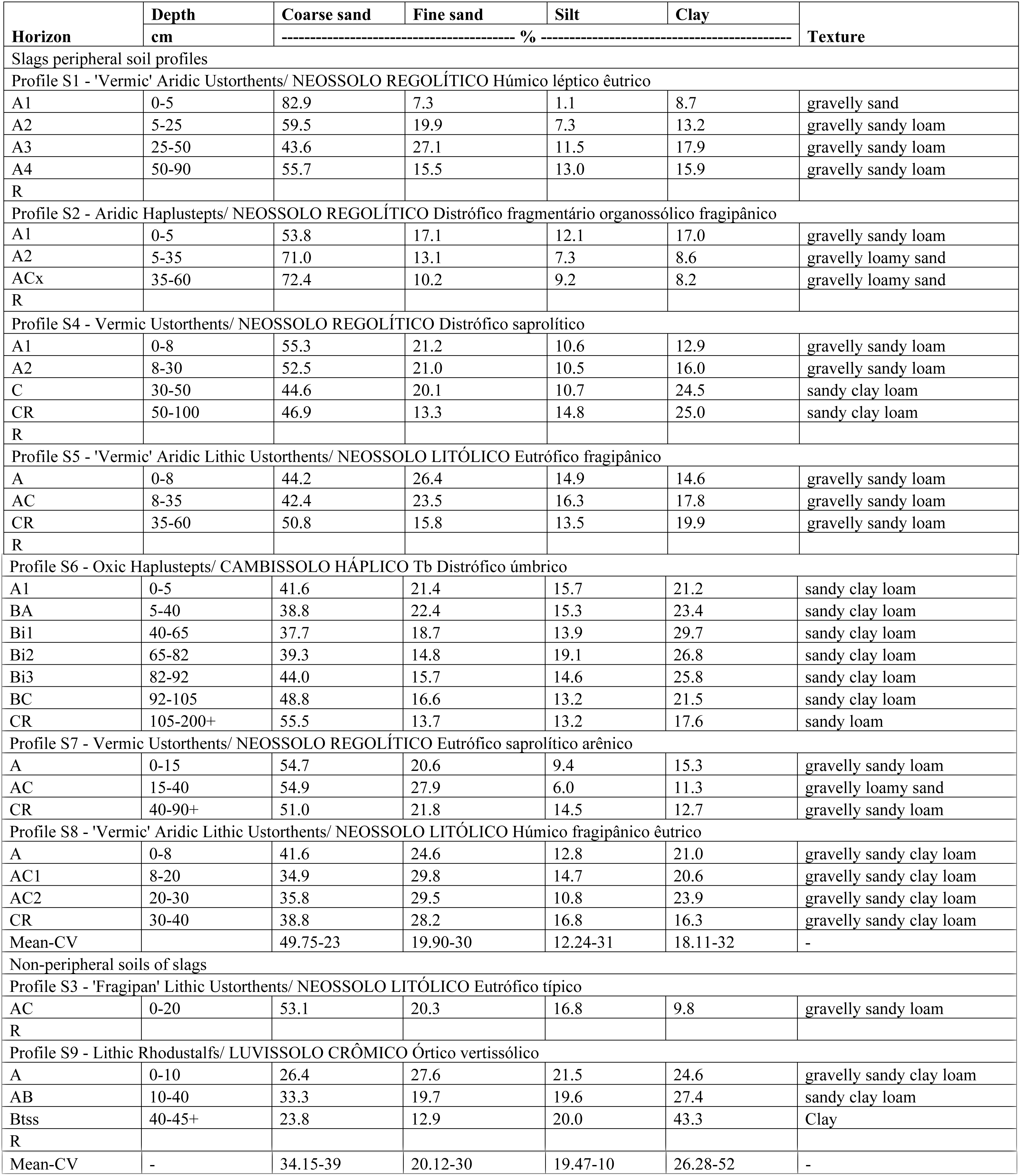
Physical soil properties.

The pedons are strongly acid to neutral and have base saturation (V) above 50 % in all horizons, except Oxic Haplustepts (Table 5). Ca^2+^>Mg^2+^>K^+^>Na^+^ is the base dominance in the exchange complex. High variation of Mehlich-1 extractable P contents (P_M_), sum of bases (SB), cation exchange capacity (ECEC and CEC), aluminum saturation (m), sodium saturation (ISNA), total organic carbon (C), total nitrogen (N) and carbon to nitrogen ratio (C/N) were detected. KCl pH is equal or below 5.0 and delta pH is negative in all horizons, indicating that, even the pedons with the lowest total organic carbon content, have dominance of negative charges.

**Table 5.**
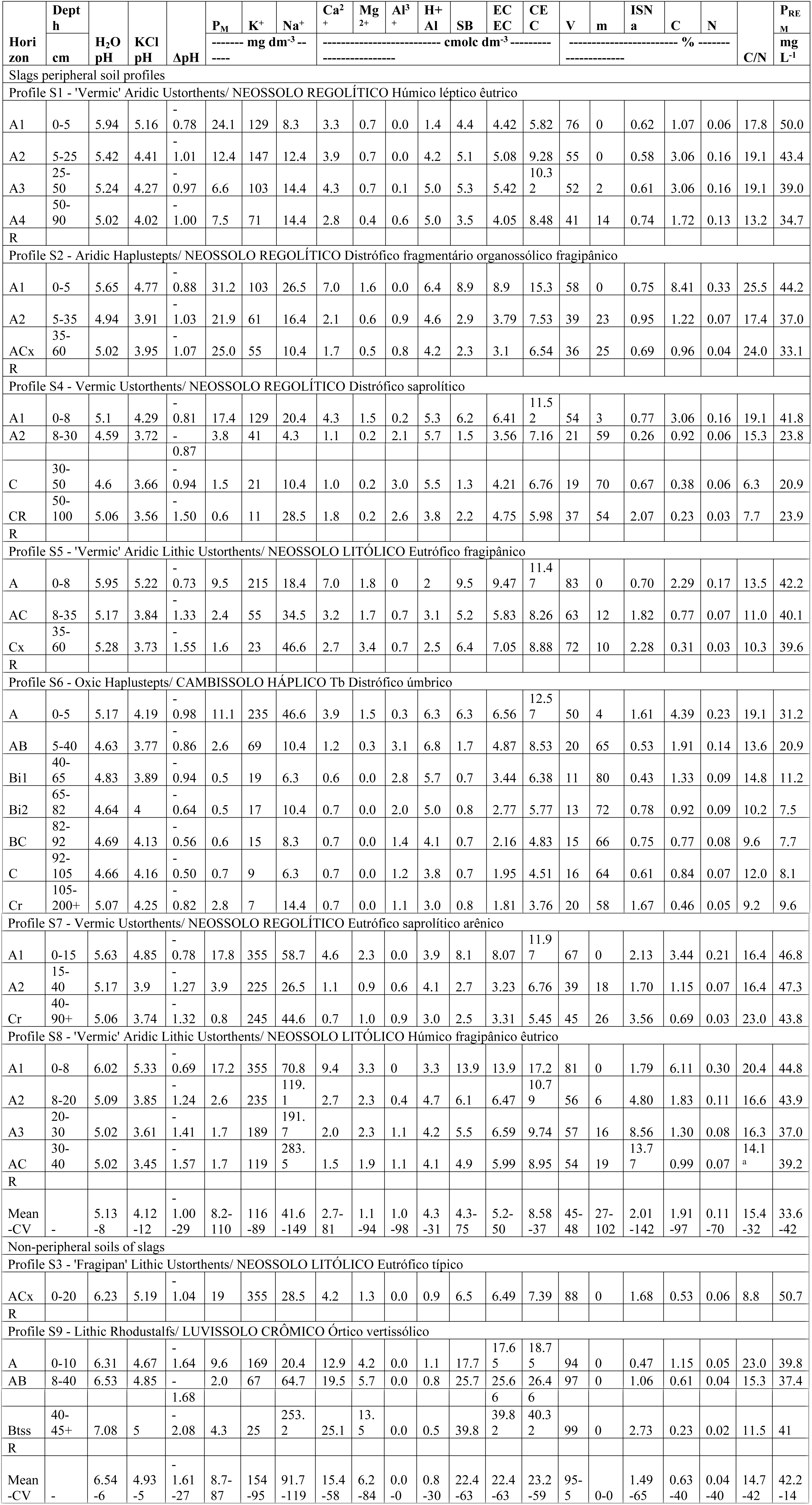
Chemical soil properties of massive influenced and non-influenced soils.

Remaining P (P_REM_) is dominantly above 30 mg L^-1^ and low variable. The higher the value of P_REM_, the lower is the affinity of soils for the P in the solution. High values of P_REM_ in all pedons, except by Oxic Haplustepts, indicates mineralogy dominantly silicate, and that iron oxides formation and brunification are limited.

Total organic carbon, N and C/N decrease with increase of depth. The C content in the upper horizon ranged between 3.6 ± 2.5 % (peripheral slags pedons) and 0.8 ± 0.4% (non-peripheral pedons of slags). In the 10–40 cm increment, the C content ranged from 1.8 ± 0.9% (peripheral slags pedons) to 0.4 ± 2.5% (non-peripheral pedons of slags). The mean C/N in the uppermost horizons ranges between 16.5 ± 3.7 (peripheral slags pedons) and 15.9 ± 10.0 (non-peripheral pedons of slags). In the 10–40 cm increment, the values ranged between 16.1 ±0.04 (peripheral slags pedons) and 13.4 ±13.8 (non-peripheral pedons of slags). Low correlation coefficient values indicated that P_M_, SB, CEC and V are poorly correlated to C, N and C/N (Fig 3).

**Fig 3.**
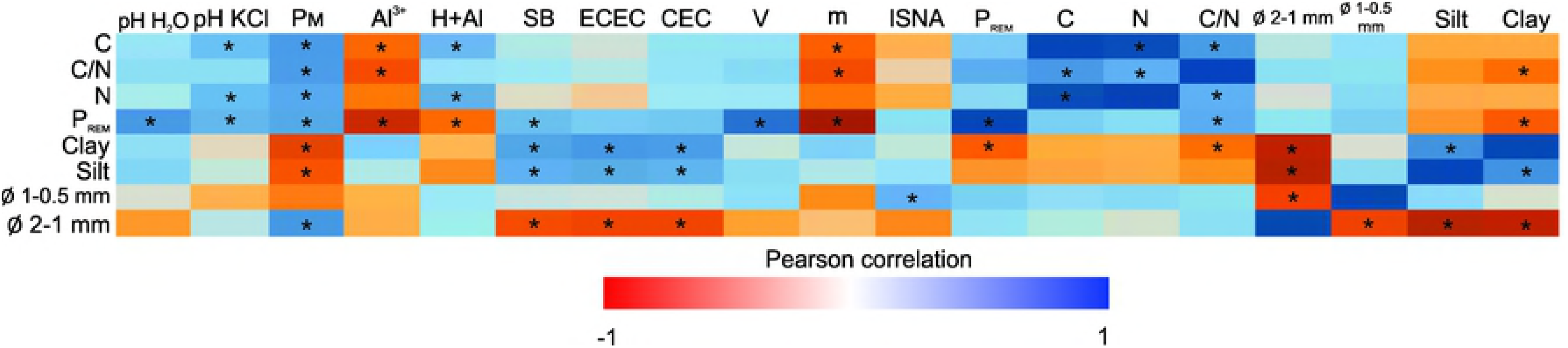
Correlation matrix between soil properties.

Both peripheral and non-peripheral soils of slags presented median values of particle density close to density of quartz (2.5-2.8 kg dm^-3^) (Table 6). Peripheral and non-peripheral soils of slags have median values of bulk density categorized as humic and sandy soils, respectively. Total porosity, macroposity, total organic carbon stock and total nitrogen stock in non-peripheral soils of slags are 16 %, 30 %, 69 % and 40 % lower compared to peripheral slags soils, respectively.

**Table 6.**
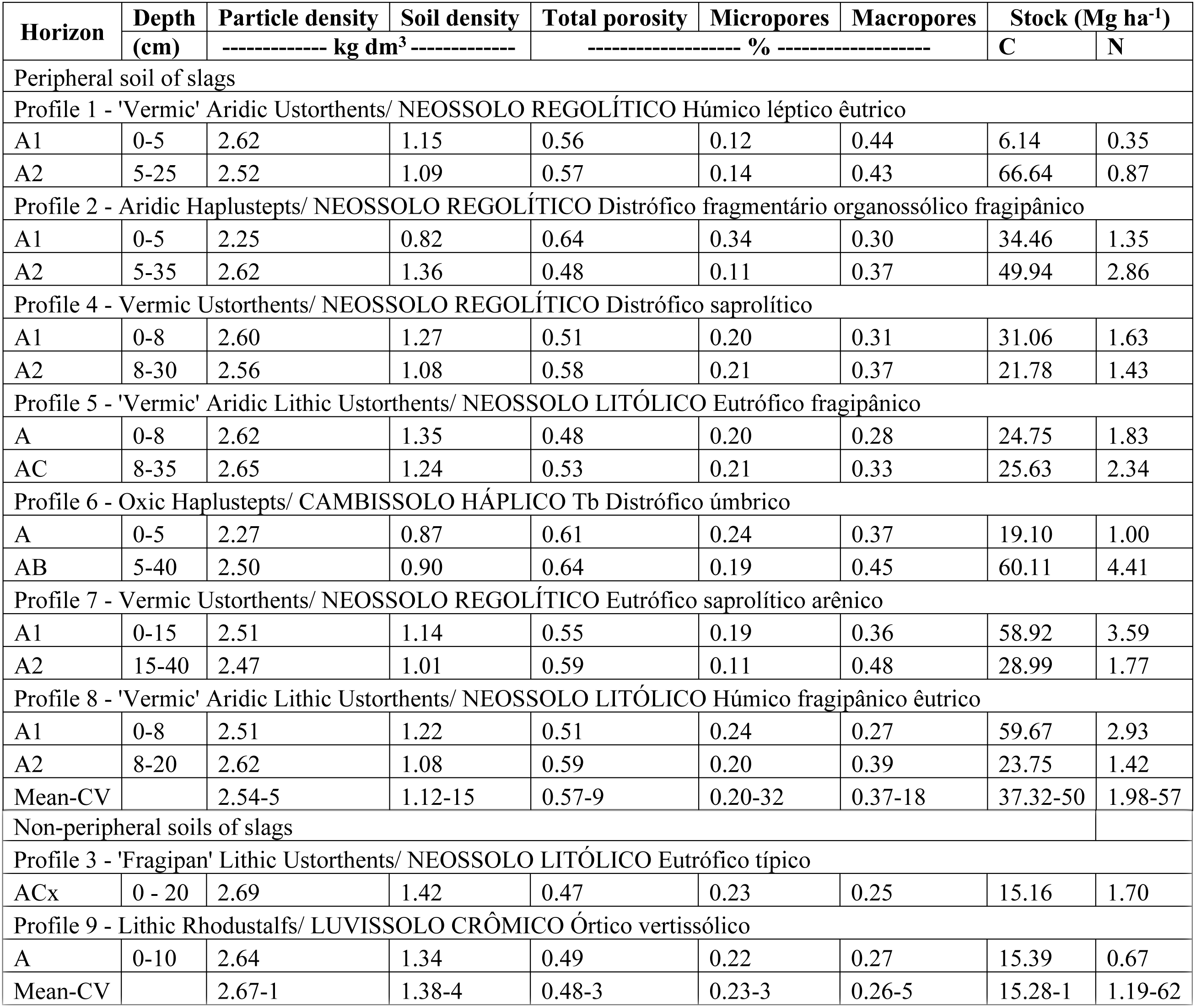
Physical soil properties determined by undisturbed samples and total organic carbon and total nitrogen stocks.

| Horizon | Depth | Particle density | Soil density | Total porosity | Micropores | Macropores | Stock (Mg ha<sup>-1</sup>) |  |
| --- | --- | --- | --- | --- | --- | --- | --- | --- |
|  | (cm) | ----- kg dm<sup>3</sup> ----- |  | ----- % ----- |  |  | C | N |
| Peripheral soil of slags |  |  |  |  |  |  |  |  |
| Profile 1 - 'Vermic' Aridic Ustorthents/ NEOSSOLO REGOLÍTICO Húmico léptico êutrico |  |  |  |  |  |  |  |  |
| A1 | 0-5 | 2.62 | 1.15 | 0.56 | 0.12 | 0.44 | 6.14 | 0.35 |
| A2 | 5-25 | 2.52 | 1.09 | 0.57 | 0.14 | 0.43 | 66.64 | 0.87 |
| Profile 2 - Aridic Haplustepts/ NEOSSOLO REGOLÍTICO Distrófico fragmentário organossólico fragipânico |  |  |  |  |  |  |  |  |
| A1 | 0-5 | 2.25 | 0.82 | 0.64 | 0.34 | 0.30 | 34.46 | 1.35 |
| A2 | 5-35 | 2.62 | 1.36 | 0.48 | 0.11 | 0.37 | 49.94 | 2.86 |
| Profile 4 - Vermic Ustorthents/ NEOSSOLO REGOLÍTICO Distrófico saprolítico |  |  |  |  |  |  |  |  |
| A1 | 0-8 | 2.60 | 1.27 | 0.51 | 0.20 | 0.31 | 31.06 | 1.63 |
| A2 | 8-30 | 2.56 | 1.08 | 0.58 | 0.21 | 0.37 | 21.78 | 1.43 |
| Profile 5 - 'Vermic' Aridic Lithic Ustorthents/ NEOSSOLO LITÓLICO Eutrófico fragipânico |  |  |  |  |  |  |  |  |
| A | 0-8 | 2.62 | 1.35 | 0.48 | 0.20 | 0.28 | 24.75 | 1.83 |
| AC | 8-35 | 2.65 | 1.24 | 0.53 | 0.21 | 0.33 | 25.63 | 2.34 |
| Profile 6 - Oxic Haplustepts/ CAMBISSOLO HÁPLICO Tb Distrófico úmbrico |  |  |  |  |  |  |  |  |
| A | 0-5 | 2.27 | 0.87 | 0.61 | 0.24 | 0.37 | 19.10 | 1.00 |
| AB | 5-40 | 2.50 | 0.90 | 0.64 | 0.19 | 0.45 | 60.11 | 4.41 |
| Profile 7 - Vermic Ustorthents/ NEOSSOLO REGOLÍTICO Eutrófico saprolítico arênico |  |  |  |  |  |  |  |  |
| A1 | 0-15 | 2.51 | 1.14 | 0.55 | 0.19 | 0.36 | 58.92 | 3.59 |
| A2 | 15-40 | 2.47 | 1.01 | 0.59 | 0.11 | 0.48 | 28.99 | 1.77 |
| Profile 8 - 'Vermic' Aridic Lithic Ustorthents/ NEOSSOLO LITÓLICO Húmico fragipânico êutrico |  |  |  |  |  |  |  |  |
| A1 | 0-8 | 2.51 | 1.22 | 0.51 | 0.24 | 0.27 | 59.67 | 2.93 |
| A2 | 8-20 | 2.62 | 1.08 | 0.59 | 0.20 | 0.39 | 23.75 | 1.42 |
| Mean-CV |  | 2.54-5 | 1.12-15 | 0.57-9 | 0.20-32 | 0.37-18 | 37.32-50 | 1.98-57 |
| Non-peripheral soils of slags |  |  |  |  |  |  |  |  |
| Profile 3 - 'Fragipan' Lithic Ustorthents/ NEOSSOLO LITÓLICO Eutrófico típico |  |  |  |  |  |  |  |  |
| ACx | 0 - 20 | 2.69 | 1.42 | 0.47 | 0.23 | 0.25 | 15.16 | 1.70 |
| Profile 9 - Lithic Rhodustalfs/ LUVISSOLO CRÔMICO Órtico vertissólico |  |  |  |  |  |  |  |  |
| A | 0-10 | 2.64 | 1.34 | 0.49 | 0.22 | 0.27 | 15.39 | 0.67 |
| Mean-CV |  | 2.67-1 | 1.38-4 | 0.48-3 | 0.23-3 | 0.26-5 | 15.28-1 | 1.19-62 |

Two main components (PCs) presented eigenvalues above 1.0, and together they explained 72.3 % of the variance (Fig 4). Component 3 and higher had eigenvalues below 1.0 and hardly explained the variance better than the original variables (1/12 = 8.3 %), so they were excluded from subsequent analyzes [35,37].

**Fig 4.**
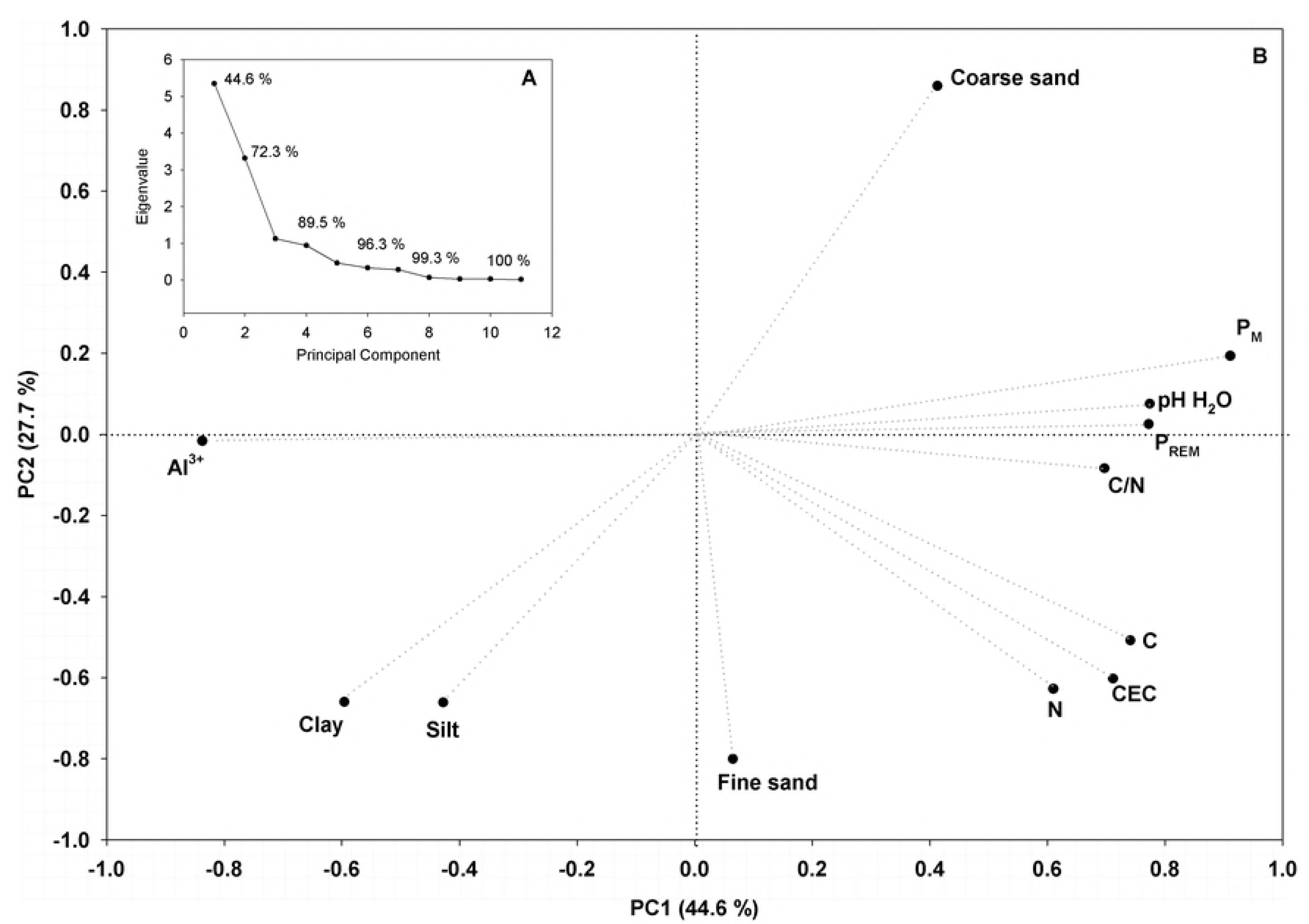
Eigenvalues of correlation matrix and explained variance per PC (A). Projection of the variables on factor plane PC1 × PC2 (B).

The first component (PC1) explained 44.6 % of the total data variance. PC1 presented high negative charge of Al^3+^ and positive charges of P_M_, C, N, C/N, and P_REM_ (Fig 4). The second component (PC2) accounted for 27.7 % of the total variance and presented large loads of CEC, coarse and fine sand, clay, silt contents. The antagonism between clay, silt, coarse and fine sand shown by PC2 could be associated to weathering process.

## Discussion

Abundance of weatherable minerals in the coarse fraction, a sandy loam texture, shallow soils, high base saturation and fragipan indicates an incipient stage of pedogenesis compared to other biomes (Tables 4 and 5). Widespread occurrence of fragipan is attributed to the restricted removal of silica in semiarid climate, which favors fragipan formation in deeper horizons [38,39]. Low sodium saturation (ISNA) indicates that salinization and sodification did not occur in minimally disturbed areas, even in the driest region of Brazil.

Poorly known, soils in caatinga are described as monotonous and low variable [40]. However, differences between peripheral slags and non-peripheral slags soils in semiarid climate are notable. Soils in the periphery of slags are deeper, has a higher number of horizons, has aggregates more developed, has lower chroma and values in topsoil, has higher total organic carbon and total nitrogen contents, and are more porous. Low value and chroma in peripheral slags soils is attributed to a higher supply of organic matter, probably related to the denser vegetation cover, higher C and N contents and C/N. These soils also presented granular structure, with larger and more frequently pores derived from intense fauna activity [41,42]. In tropical wet areas, bioturbation is related to well-structured soils through the increase of clay, carbon and nutrient contents and soil porosity [43–46].

Notification of Bi horizon is unexpected to Brazilian semiarid climate, once Inceptsols are rare in soil maps [47,48]. A harder, firmer, more sticky and plastic consistence indicates more developed horizons. Development of Inceptsol seems to be drastically related to local geomorphology, once it covers a concave depression which accumulates water and sediments percolated on slags.

The absence of coatings in the argic horizon of Lithic Rhodustalfs can be attributed to neoformation of clays or elutriation, despite eluviation [49]. The eluviation can be defined as the translocation of soil particles from surface to deeper horizons. Once eluviation depends on vertical water movement, it occurs when precipitation exceeds evaporation. Occurrence of slickensides in the argic horizon indicate marked changes in moisture content [50].

Total porosity is composed by macro and microporosity. Micropores are formed by congruent dissolution of minerals and soil biota and are usually found within structural aggregates. The water within these pores is considered immobile, but available for plant extraction. So, these smaller pores are important to retain water required for plant during dry season. The lower values of microporosity in peripheral soils of slags suggest that the high vegetable cover explores water from other sources, as litter [51] or atmosphere [52]. On the other hand, macroporisity is a highly dynamic soil property once it is influenced by environmental factors, as roots, soil biota and raindrop impact, wetting-drying cycles and human activities [53–55]. Low difference between macroporosity of surface horizon and its underlying guarantees water stocking by vertical water flux from surface to deeper horizons. The high density of roots close to lithic contact and fragipan suggests that water did not percolate below the active root zone by the force of gravity. Furthermore, sandy loam texture dominance reduces loss of water by capillarity-evaporation from soil surfaces [56].

Mean total organic carbon (C) is below other Brazilian biomes [57–60] and mean C/N is until 1.7 times lower than Amazonian rainforest [61]. The amount of C is determined by the net balance between the rate of input as leaf and root biomass and its mineralization. Soil structure is a key element in C dynamic. The increase of silt and clay contents during weathering, and its organization in aggregates, protects organic carbon from oxidation [62,63]. In dry tropical climates, carbon stock in the upper-most 30 cm of mineral soils estimate ranges from 31 Mg C ha^-1^ in sandy soils to 38 Mg C ha^-1^ in high activity clay soils [64]. However, the weak correlation between texture and C (Fig 3) suggest that stabilization of organic matter by the formation of stable complexes with clay minerals did not occurs or it is underestimated due to sand dominance.

Weak correlation between total organic carbon and CEC or base saturation indicates that soil fertility is not associated to total organic matter and, possibly related to parent material. This result indicates that paludization is uncapable to influence geochemical processes in caatinga, in opposite to other biomes [65–68].

C/N also determines total organic matter decomposition. A decrease in C/N simultaneous to the proportion of stabilized organic carbon compounds has been shown in several studies [69–71]. Strong positive correlation between C/N and C indicates that the organic carbon is stocked in less favorable conditions for microbial activity. Higher total organic carbon in peripheral slags soils highlights massive as crucial areas for the management of C sequestration in Caatinga.

The mean total organic carbon stock was 35 % higher than previously studies in Caatinga [72]. Soil organic carbon stock is commonly related to clay content and altitude [72], water availability and soil organic matter content in Caatinga [73]. Here, we observed that local geomorphology is more important to carbon sequestration in Caatinga, once massive influenced areas are larger than highlands. In opposite to previous studies [74], we did not observed a predominant decrease of total organic carbon and total nitrogen stocks with increasing of depth. This result can be attributed to several causes: a) total organic carbon and nitrogen are higher in denser vegetable cover V.P.; b) total organic carbon content is defined by an equilibrium between the low organic residue inputs [75] and high C/N values (Table 5); c) virtual absence of clay- organic complexes evidenced by low correlation between clay content and soil organic carbon (Fig 3); d) absence of eluviation evidences suggest that organic complexes are not transported by water vertical movement (Table 3).

Low similarity of plant communities between plots on peripheral areas of slags and plots on a flat plain (Fig 2) suggest that soil is the main limiting factor for phytogeographical patterns [76–79]. High tree species richness can be related to low levels of physiological stress, product of sites with high soil fertility [10,80]. Previous studies suggested that the same pattern in Caatinga [81], however the relative low coefficient of variation of base saturation suggest that shrub-arboreal diversity and abundance could be related to other soil properties.

The higher number of specimens in slags peripheral soils (S1 Table), especially of these species documented in (sub)humid biomes, is attributed to increase of water retention due to high soil organic carbon, a valuable resource in xeric shrubland. Root depth in massive influenced pedons were higher than previously studies in Caatinga soils [82]. So, we believe that trees exploit a larger volume in peripheral slags soils, which guarantees them mechanisms of resistance to the climatic seasonality and consequently diversify the environments [83]. Although the proportion of humin, fluvic acids and humic acids influence the relation between water retention and soil organic matter content, a 1 % mass increase in soil organic matter content, on average, increases water content at saturation, field capacity, wilting point and available water capacity by: 2.95, 1.61, 0.17 and 1.16 mm H_2_O 100 mm soil^−1^, respectively [84]. The increase is larger in sandy soils, followed by loams and is least in clays [85].

These narrow areas around the slabs present floristic similarities with some tropical forests existing in highlands environments in the Brazilian semiarid [24,86]. This study reveals that cover of plant species considered exclusive of (sub)humid biomes in Brazil extends beyond highlands in the semiarid, associated with the high soil organic carbon content and water retention capacity of more developed soils than the typical of the Caatinga. These results corroborate the hypothesis that diversification of niches favors increase of plant species diversity in tropics [87].

However, if maintenance of this pedological and botanical conditions in a predominantly semiarid climate seems reasonably deciphered, questions arise about how high water-demand species reached semiarid regions. We believe that vegetable cover in areas influenced by slags and massives in the Caatinga are testimony to an old existing vegetable cover, inherited from a more humid past climate, and confined to refuges with the establishment of the current semiarid climate. Such vegetable cover would have existed between Late Miocene and Early Pleistocene, uniting the current Amazon, Atlantic and Caatinga biomes [6, 88–92]. European colonization in the sixteenth century and the advance of deforestation since then also could have contributed to the fragmentation of refugees [93].

Peripheral areas of slags are important today and can be decisive in future. Eighty percent of total transpiration in Caatinga is controlled by top layers of soil (0 – 20 cm) [94]. According to prevision of a rise in temperature between 0.5 and 4° C and a reduction of precipitation of 10–20 % for the Brazilian Northeast region by 2,100 [95] the Caatinga biome may become completely soil-water pulse dominated. Consequently, soil organic carbon stock also could be reduced by decrease of organic residue input. Future studies should investigate biotic responses from these refuges to climate changes [96].

## Conclusions

Despite the semiarid surrounding context, pedogenetic processes typical of tropical wet conditions, such as paludization, melanization and bioturbation were verified in slags peripherical areas. Well-developed soils with high soil organic matter and water holding capacity support semi-deciduous forests of (sub)humid biomes. These areas constitute a refuge for several plant species that cannot develop on the typical soils of the semiarid. Furthermore, future vegetation surveys should focus different habitats to expand plant diversity knowledge of the Caatinga.

Carbon stock data is rare and soil diversity is poorly known in Caatinga. Further research could increase estimates of semiarid soil carbon stocks by 37 to 69 %, helping to reduce uncertainty about the global carbon budget. Our findings could also lead to the development of innovative conservation and land restoration actions in dryland biomes regions with low opportunity cost to mitigate climate change, combat desertification, and support conservation of biodiversity and ecosystem services that underpin human livelihoods.

## Supporting information

**S1 Table. Number of specimens identified in vegetational plots by family, scientific name and its reported biomes according to literature.**

